# Correlated evolution of conspicuous coloration and burrowing in crayfish

**DOI:** 10.1101/2023.07.03.547601

**Authors:** Zackary A. Graham, Dylan J. Padilla Perez

## Abstract

Conspicuous colors have fascinated biologists for centuries, leading to much research on the evolution and functional significance of color traits. However, some authors have critiqued the adaptationist dogma amongst color researchers. When investigating a color trait, researchers often exclusively consider the alternative hypotheses—they assume color is adaptive. The null hypothesis of animal color—that coloration is non-adaptive or evolutionary neutral, is rarely considered. Here, we use phylogenetic comparative methods to investigate color evolution throughout freshwater crayfishes. Within the taxa we analyzed, conspicuous colors have evolved independently over 50 times. The intuitive, but not evolutionary-justified assumption when presented these results is to assume that these colors are an adaptation. But contrary to this intuition, our work might support the hypothesis that coloration in crayfish is neutral; because we show that conspicuous colors are evolutionary correlated to a semi-terrestrial burrowing lifestyle. Conspicuous coloration being common in semi-terrestrial burrowers is paradoxical, because these species are nocturnal, and rarely leave their burrows. Overall, our work brings into question to traditional view of animal coloration as a perfectly adapted phenotype.

## Introduction

Animals with bright, conspicuous colors fascinate evolutionary biologists and the public (Cuthill et al., 2017). Unlike cryptic colors, which minimize detection, conspicuous colors are unlikely to appear in the natural environment and draw attention to an animal (Stevens and Merilaita, 2011; Ruxton et al., 2018; Emberts and Wiens, 2022). The functional significance of conspicuous colors has been the focus of more than two centuries of evolutionary thought, with both Charles Darwin and Alfred Russel Wallace using natural selection to explore and explain the utility of puzzlingly conspicuous colors such as the peacocks plumage or the contrasting color patterns of elapid snakes (Wallace, 1877; Darwin, 1888; Caro, 2017). Although the evolution of conspicuous colors was once considered an evolutionary paradox (Marples et al., 2005; Briolat et al., 2019), the functional significance of these colors are now clear in many taxa. Warning coloration and sexual selection have been the dominant theoretical forces that explain the evolution of conspicuous colors (Ruxton et al., 2018; Emberts and Wiens, 2022; Andersson, 1994). As such, conspicuous colors are thought to be adaptive if they communicate information to receivers, whether it be unprofitable consumption (i.e., warning coloration including aposematism, long handling times, etc.) or some index of quality (i.e., sexual selection through female mate choice or male-male competition). Indeed, famous examples of conspicuously colored animals can be at least partially explained by one of these two mechanisms, like warning coloration in poison dart frogs (Hagman and Forsman, 2003; Maan and Cummings, 2012), hymenopterans (Chatelain et al., 2023), and coral snakes (Davis Rabosky et al., 2016). And sexually selected colors in birds (Dunn et al., 2015; Weaver et al., 2017), tropical freshwater fishes (Endler, 1984; Maan and Sefc, 2013), and dragonflies (Suárez-Tovar et al., 2022). In addition to warning coloration and sexual selection, alternative functions such as species identification or crypsis may explain the functional significance of conspicuous colors (Losos, 1985; Johnsen, 2005; Wollenberg and John Measey, 2009; Perez and Backwell, 2020).

Our understanding of the proximate mechanisms that generate the diversity of conspicuous colors has advanced significantly with the advent of modern technology. Genetic, physiological, and structural underpinnings of colors are now widely studied, especially in vertebrates like birds, mammals, and lizards (Hill, 1996; Rosenblum et al., 2004; Hoekstra et al., 2006; Walsh et al., 2012; Maan and Sefc, 2013; Cuthill et al., 2017). By contrast, comparative studies on the ultimate mechanisms that select for conspicuous coloration are scarce. For example, although contemporary biologists acknowledge the costs and benefits of conspicuous coloration, less is known about what drives one species to be conspicuously colored, but closely related species to remain cryptically colored. Recent studies have shed light on this question. Emberts and Wiens (2022) found that diel patterns predict the evolution and function of conspicuous coloration across land vertebrates. In their analyses, warning coloration was more likely to evolve in groups that are nocturnal (or ancestrally nocturnal) and sexually dichromatic colors being more likely to evolve in diurnal species (or ancestral diurnal). These results are intuitive, as the light environment during activity (i.e., being active diurnally or nocturnally) strongly relates to animal coloration (Endler, 1984; Warrant and Johnsen, 2013). Predation is likely to be another strong predictor as to whether conspicuous colors are adaptive or maladaptive. Indeed, in poison dart frogs, conspicuous warning coloration may be evolutionary correlated to body size, such that visually guided predators select for the evolution of larger body sizes in species with conspicuous colors or vice versa (Hagman and Forsman, 2003). A species life and natural history may similarly influence the evolution of conspicuous colors. In birds, both habitat type and mating system can influence whether a species is cryptic, conspicuous, and the degree of sexual dichromatism (Dunn et al., 2015). These studies demonstrate a few variables known to influence an animal’s light environment, which in turn alters the costs and benefits of conspicuous versus cryptic coloration.

Despite recent theoretical and empirical studies demonstrating the evolutionary drivers of conspicuous coloration, biologists still struggle to explain the functional significance of color in some animals. One such case is cave-dwelling or fossorial animals which spend most of their lives hidden from daylight (Gross and Wilkens, 2013). When light levels are low or non-existent, selection for protective coloration will be minimal, which leads to animals poorly matching their background. Despite this, in some cases, animals that rarely interact with the surface exhibit conspicuous coloration, presenting a paradox. In caecilians, despite this group living in soils and leaf litter, many caecilians have yellow, red, or blue dorsal body coloration (Wollenberg and John Measey, 2009). Phylogenetic analysis in this group demonstrates that coloration has evolved multiple times, leading the authors to speculate on the functional significance of their coloration. Interestingly, one proposed hypothesis suggests that caecilian coloration is the product of neutral evolution or caused by physiological/genetic constraints. However, in their study, the authors conclude that since yellow coloration is correlated with their degree of surface movements, caecilian coloration may serve dual functions in aposematism and in crypsis, which allows these animals to remain defended during rare times of surface movement (Wollenberg and John Measey, 2009). But support for this hypothesis is untested and remains speculative.

Another example in which biologists face difficulties explaining the functional significance of conspicuous coloration comes from the coconut crab, *Birgus latro* (Nokelainen et al., 2018). Coconut crabs are nocturnal and exhibit a discrete phenotypic color polymorphism with some individuals exhibiting red coloration and other conspecifics exhibiting blue coloration, typically in a 3:1 red-blue ratio (Nokelainen et al., 2018). Multiple studies have attempted to reveal the functional significance of color polymorphism in coconut crabs, with little support for any of the proposed hypotheses. Color polymorphism in coconut crabs is not linked to sex, size, age, morphology, or habitat use (Nokelainen et al., 2018; Caro, 2017; Caro et al., 2019). This frustrating series of unsupported hypotheses has led researchers to suggest that coconut crab color may not be functionally significant at all, but rather, function as a neutral trait (Caro, 2021). Several lines of evidence support this claim, including the frequencies of red and blue colors lining up with predictions from simple mendelian genetic frequencies (3:1), as well as similar color mutations which result in blue and red coloration being known from other decapods. Overall, Caro (2021) highlights the overly adaptationist viewpoint of color researchers. In this way, not all colors are predicted to perfectly match their background or be considered an adaptation. Some colors may not be under selection, but rather, they could be evolutionary by-products of mutations subject to little or no selection (Wollenberg and John Measey, 2009; Ho et al., 2017; Caro, 2021). This provocative, but potentially informative neutral theory of color evolution reviewed by Caro (2021) brings a nuanced view to biologists’ understanding of animal color. However, the hypothesis that coloration may evolve as a neutral trait remains to be assessed in a robust comparative framework.

In the current study, we use phylogenetic comparative methods to test the hypothesis that conspicuous coloration can evolve and be maintained as an evolutionary neutral trait. We do so in freshwater crayfish (Decapoda: Astacidea), which are an ideal group for this test for several reasons. First, although crayfish are most often associated with cryptic colors that aid in camouflage from their primarily visually-oriented predators, with over 700 species, crayfish possess a range of conspicuous colors and color patterns—with many species being blue, red, orange, purple, or a combination of these colors (Schuster, 2020, Figure 1). Intriguingly, there is little to no evidence supporting the functional significance of conspicuously colored crayfish, making this an evolutionary puzzle (Schuster, 2020). Second, crayfish vary on a spectrum of burrowing and surface activity, which allows us to evaluate our hypothesis in species with divergent degrees of predation pressure, migration, and gene flow Hobbs Jr (1981); Hurt et al. (2019). About 70% of crayfish species live in permanent bodies of water and construct rudimentary burrows within the benthic and hyporheic zones (i.e., aquatic burrowing crayfish). However, a semi-terrestrial burrowing lifestyle has evolved over a dozen times throughout the crayfish phylogeny (Breinholt et al., 2012; Stern et al., 2017). Semi-terrestrial crayfish burrows can reach several meters down to the water table, with multiple surface entrances and underground chambers (Richardson, 2007). Most semi-terrestrial burrowing crayfish have small geographic ranges due to poor dispersal abilities over land, resulting in lower population sizes, genetic diversity, and gene flow compared to aquatic burrowing crayfish (Ponniah and Hughes, 2004; Stern et al., 2017; Hurt et al., 2019). Moreover, semi-terrestrial burrowing crayfishes are also thought to be more likely to be conspicuously colored, but no systematic study has verified these claims (Schuster, 2020).

**Figure 1:**
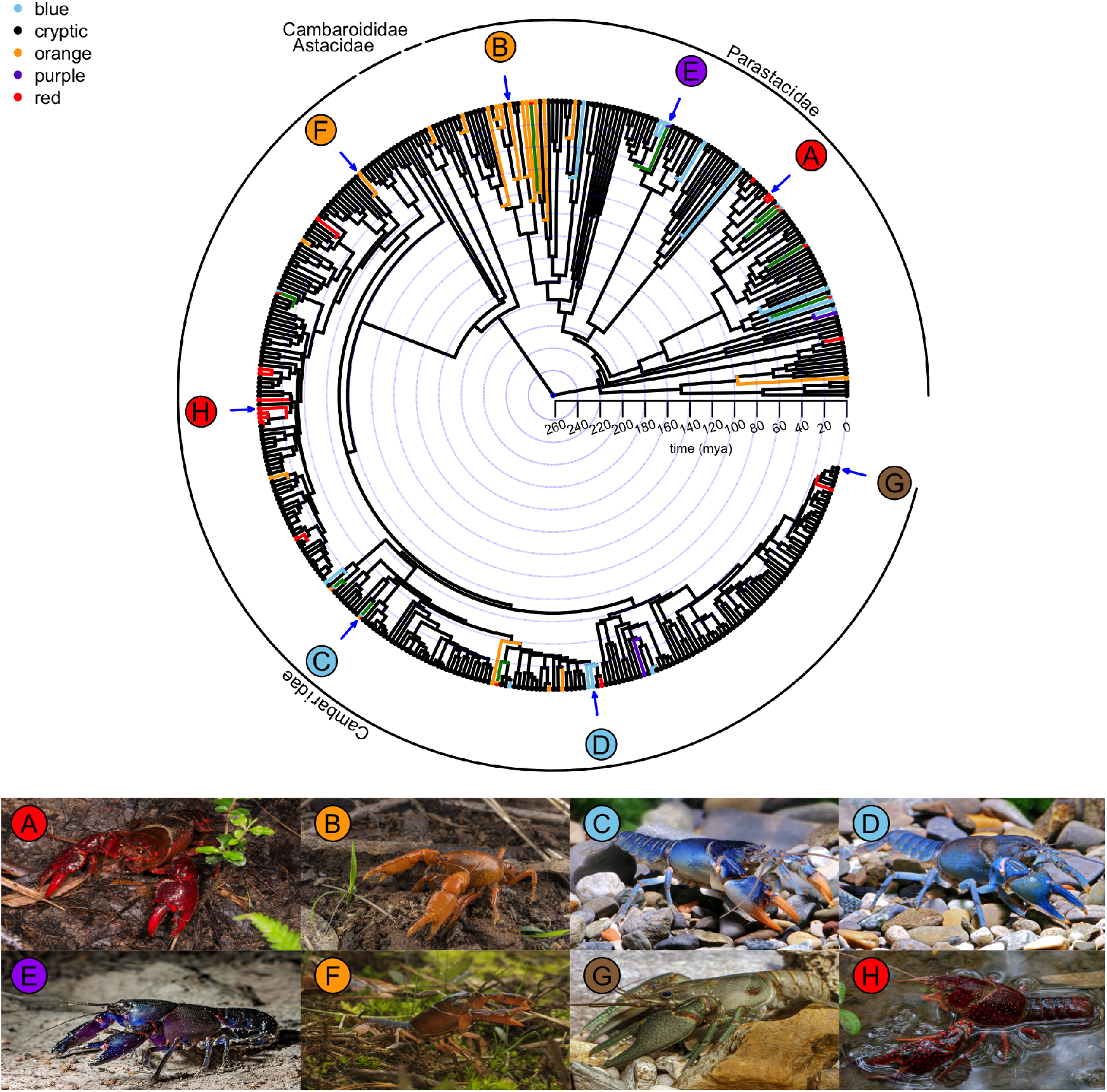
Evolution of coloration throughout 406 species of freshwater crayfish. Branch lengths and branch tips are colored based on the species that exhibit cryptic and conspicuous coloration. Green branches represent polymorphic species. Photographs highlight the diversity of colors in crayfish species; A) *Euastacus astralasiensis*, B) *Engaeus cymus*, C) *Cambarus dubius*, D) *Cambarus harti*, E) *Cherax robustus*, F) *Distocambarus carlsoni*, G) *Faxonius rusticus*, H) *Procambarus clarkii*. Photo credits: A) Ciaran Nagle, B) Ryan Francis, C) and D) Chris Lukhaup, E) Narelle Power, G) Guenter Schuster, H) fra298 on Flickr.

Conspicuous coloration being more common in semi-terrestrial burrowers is paradoxical, because these species are primarily nocturnal, and spend the majority of their lives hidden underground (Bearden et al., 2021). Because genetic mechanisms such as drift and mutation events are one possible explanation of neutral color traits (Caro, 2021), this makes the contrast between crayfish burrowing strategies an ideal system to test neutral hypothesis of color evolution. In theory, similar to what has been suggested in coconut crabs (Caro, 2021); a color mutation may arise in a population of semi-terrestrial crayfish and become a unique color phenotype if there is a small population size, and minimal gene flow and migration rates (Martin and Orgogozo, 2013). Lastly, crayfish are an ideal system to investigate our hypothesis because the physiological underpinnings of coloration in decapod crustaceans is well studied. Decapod color is primarily driven by the layering of carotenoid pigments and crustacean specific carotenoprotein complexes that deposit themselves in different layers of the epithelial and cuticular tissues (Cianci et al., 2002; Tlusty and Hyland, 2005; Wade et al., 2009, 2017). Together, these factors make crayfish an ideal group to investigate the neutral theory of color evolution.

If conspicuous colors has evolved as a neutral trait in crayfish, we predict that semiterrestrial burrowing crayfishes will be more likely to exhibit conspicuous coloration compared to aquatic burrowing species. However, if we find that semi-terrestrial burrowers and aquatic burrowers are equally as likely, or that aquatic burrowers are more likely to have conspicuous colors, this suggests that some other functional mechanism may explain the evolution of coloration in crayfishes. Ultimately, we explore multiple potential hypotheses to explain the functional significance (or lack thereof) of conspicuous coloration across crayfishes and present the first phylogenetically informed analysis of coloration is this group.

## Materials and Methods

### Phylogeny

We used a recent time-calibrated phylogeny of 466 species of crayfish. This phylogeny was estimated using a maximum likelihood analysis of 1565 total sequences from 466 taxa (*≈* 60% of the described diversity). The tree was dated using penalized likelihood and eight fossil calibration points (Stern et al., 2017). We then pruned the phylogeny to the crayfish species that we had data for each analysis.

### Coding of species color

For each species within the phylogeny, we compiled information on their coloration using reports from the primary literature, secondary literature, as well as color photographs from online and print materials (Hobbs Jr, 1981; Pflieger and Dryden, 1996; Taylor and Schuster, 2004; Walls, 2009; Schuster, 2020; Schuster et al., 2022). Photographs of juvenile crayfish or crayfish that showed evidence of recent molting were discarded and not considered, because crayfish color may change throughout ontogeny or the molt cycle (Schuster, 2020). Then, each species was classified as having either cryptic or conspicuous coloration. In all cases, we only determined a crayfish to have conspicuous or cryptic coloration based on the dominant color present throughout their body. For example, many *Faxonius* spp. have black and orange contrasting bands at the tips of their claws. Other crayfish have small, highlights of red surrounding their rostrum or join appendages (i.e., *Lacunicambarus ludovicianus*). Similarly, some crayfishes may have legs that are blue-tinted (i.e., *Cambarus jezerinaci*), but the remainder of their bodies is brown. In all of these instances described above, if the dominant color of the exoskeleton was cryptically colored, we would classify these crayfishes as being cryptic, instead of conspicuous. Additionally, we only coded images that showed a clear dorsal view of the crayfish exoskeleton. Thus, even though some species may have conspicuous coloration on the ventral face of their claws (i.e., red coloration in *Pacifastacus leniusculus* or *P. fortis*), we would code these species as being cryptic, not conspicuous. Notably, several genera of crayfish possess troglobitic species, which are unpigmented. We chose to code these species as cryptic because, despite these species stand out from their background in the presence of light, their natural environment is unlit and they should not be treated as conspicuous. Lastly, in species in which the conspicuous colors were mentioned but were described as being relatively cryptic (e.g., reddish-brown, grayish-blue), these colors were coded as cryptic, and not conspicuous.

If a species was coded as conspicuous, we categorized them as being one of four conspicuous colors: red, orange, blue, or purple. Yellow is also considered a conspicuous color (Emberts and Wiens, 2022), but no crayfish exhibit conspicuous yellow coloration. Some species of crayfish are conspicuously polymorphic; a single species may occur in multiple conspicuous color morphs within and across populations (i.e., red, orange, and blue, phenotypes in *Cambarus dubius*). In these cases, we coded which conspicuous colors were present in these species, and also coded them as being polymorphic.

Importantly, cryptic and conspicuous color depends on background, light environment, and the visual system of receivers (Endler, 1984; Warrant and Johnsen, 2013). However, we believe that the colors we considered conspicuous are unlikely to appear in the natural environment and therefore unlikely to be cryptic (regardless of background). Although we recognize that many crayfish are patterned with striping, banding, or dots throughout their body which could potentially be considered conspicuous (Schuster, 2020, depending on background), the focus of the color study was coloration, not patterning, and therefore these individuals were coded as cryptic. We also acknowledge that human perception of color may present biases due to the human visual system. But analysis in birds suggests that human assessments of color strongly correlates to quantitative analysis of color (Bergeron and Fuller, 2018).

Color data are available in the GitHub repository provided below, which also includes references and coding for the species included in our analyses.

### Burrowing Classification

Historically, several sources have classified crayfishes into ecological groups based on a mixture of morphology, behavior, and burrowing abilities (Hobbs, 1942; Hobbs Jr, 1981; Horwitz and Richardson, 1986; Welch and Eversole, 2006). Many of these classifications were formalized with specific taxonomic groups in mind, (Hobbs, 1942; Hobbs Jr, 1981, i.e., was primarily used for Cambaridae), and not the entire crayfish clade. Here, we used a burrowing classification which separates crayfish based on their ability to create burrows in semi-terrestrial habitats or aquatic habitats (presented in the GitHub repository provided below). Effectively, this takes the most widely used burrowing classification formalized by Hobbs (1942); Hobbs Jr (1981) and combines what was previously three ecological groups (primary, secondary, and tertiary burrowers) into two groups: semi-terrestrial burrowers and aquatic burrowers. Aquatic burrowers inhabit permanent bodies of water and construct rudimentary burrows underneath shelters within the benthic or hyporheic zone (i.e., tertiary burrower). By contrast, semi-terrestrial burrowers can construct burrows in semi-terrestrial habitats, which are often adjacent to, but may be far from permanent bodies of water (i.e., secondary and primary burrowing crayfishes). We used a recent review of burrowing crayfish life-history studies to split crayfish into primary, secondary, or tertiary burrower species, which could then be grouped together as aquatic and semi-terrestrial burrowers (Bloomer et al., 2021).

### Statistical Analysis

To test for an evolutionary relationship between conspicuous coloration and burrowing behavior, we fitted Pagel94 models using the function *fitPagel* available in the R package **“phytools”** v.1.7-0 (Pagel, 1994; Revell, 2012). We fitted 4 different models to the data described as follows: the first model consisted of a null model or independent model under which conspicuous coloration and burrowing were assumed to evolve independently from one another along the tree. The second model consisted of a dependent model under which the rates of evolution for conspicuous coloration were allowed to depend on the burrowing state and vice versa. In the third model, we allowed the rate of evolution of conspicuous coloration to depend on the burrowing state, but not the converse. In the fourth model, however, we allowed the rate of evolution of burrowing behavior to depend on the state of conspicuous coloration, but not the converse. We then compared the models using a likelihood-test ratio through the *anova* function available in **“phytools”**, which enabled us to select the most likely model based on the Akaike Information Criterion (AIC).

To produce a visualization of our results and ensure that they are fully reproducible, we carried out all the analyses in the free software for statistical computing R v.4.2.2 (R Core Team, 2022).

## Results

According to the Akaike Information Criterion and the weight of evidence, a model wherein conspicuous coloration depends on burrowing behavior, but not the converse, fits our data better than other models (Table 1).

**Table 1:** Contrast of parameters estimated by the most likely models, according to the information-theoretic criteria of selection.

| | $\log(L)$ | d.f. | AIC | weight |
| --- | --- | --- | --- | --- |
| null | -323.170 | 4.000 | 654.341 | 0.000 |
| dependent.burrowing | -315.591 | 6.000 | 643.182 | 0.091 |
| dependent.bodycol | -313.638 | 6.000 | 639.275 | 0.645 |
| interdependent.model | -312.534 | 8.000 | 641.068 | 0.263 |

If conspicuous coloration and burrowing behavior evolved independently from one another, as expected by the null model, the transition rate from conspicuous coloration to cryptic coloration would be the highest (Figure 3a). A relatively high transition rate from aquatic burrowing to semi-terrestrial burrowing can also be observed, but the difference with the converse transition rate was negligible (Figure 3a). Unsurprisingly, our data indicated that cryptic coloration is the predominant phenotype across crayfish lineages. However, the frequency of orange and blue colorations appears to be more common in semi-terrestrial burrowers. By contrast, red coloration appears to be more common in aquatic burrowers (Figure 2).

**Figure 2:**
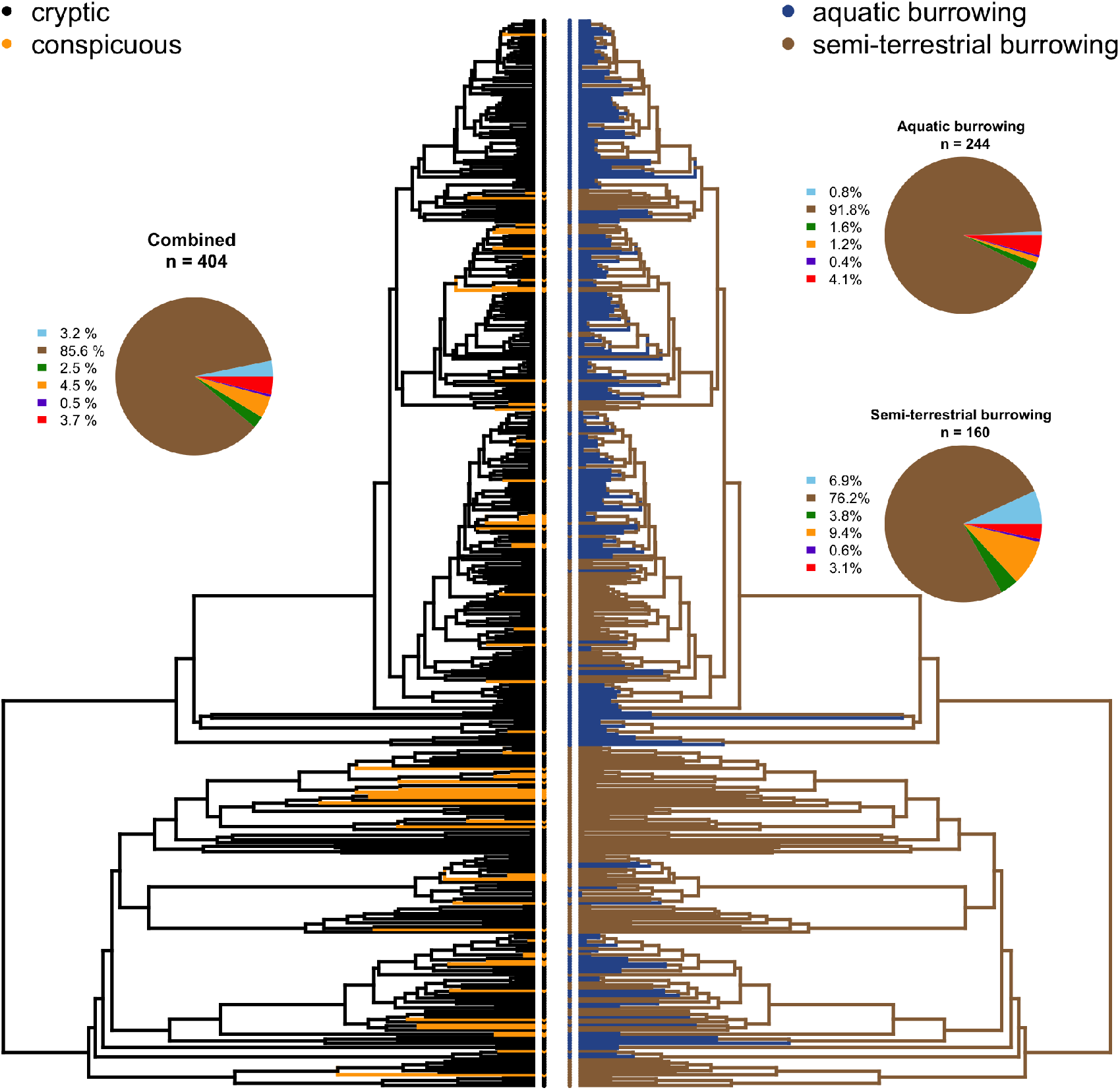
Correlated evolution of conspicuous coloration and burrowing throughout 406 species of freshwater crayfish. The left phylogeny tree depicts the lineages in which transitions from cryptic coloration to conspicuous coloration, and vice versa, have occurred. Likewise, the right phylogeny depicts the transitions from aquatic burrowing to semi-terrestrial burrowing, and vice versa. Inset pie charts show the frequency of cryptic or conspicuous colors for all species, and the frequency of aquatic vs. semi-terrestrial burrowing species.

**Figure 3:**
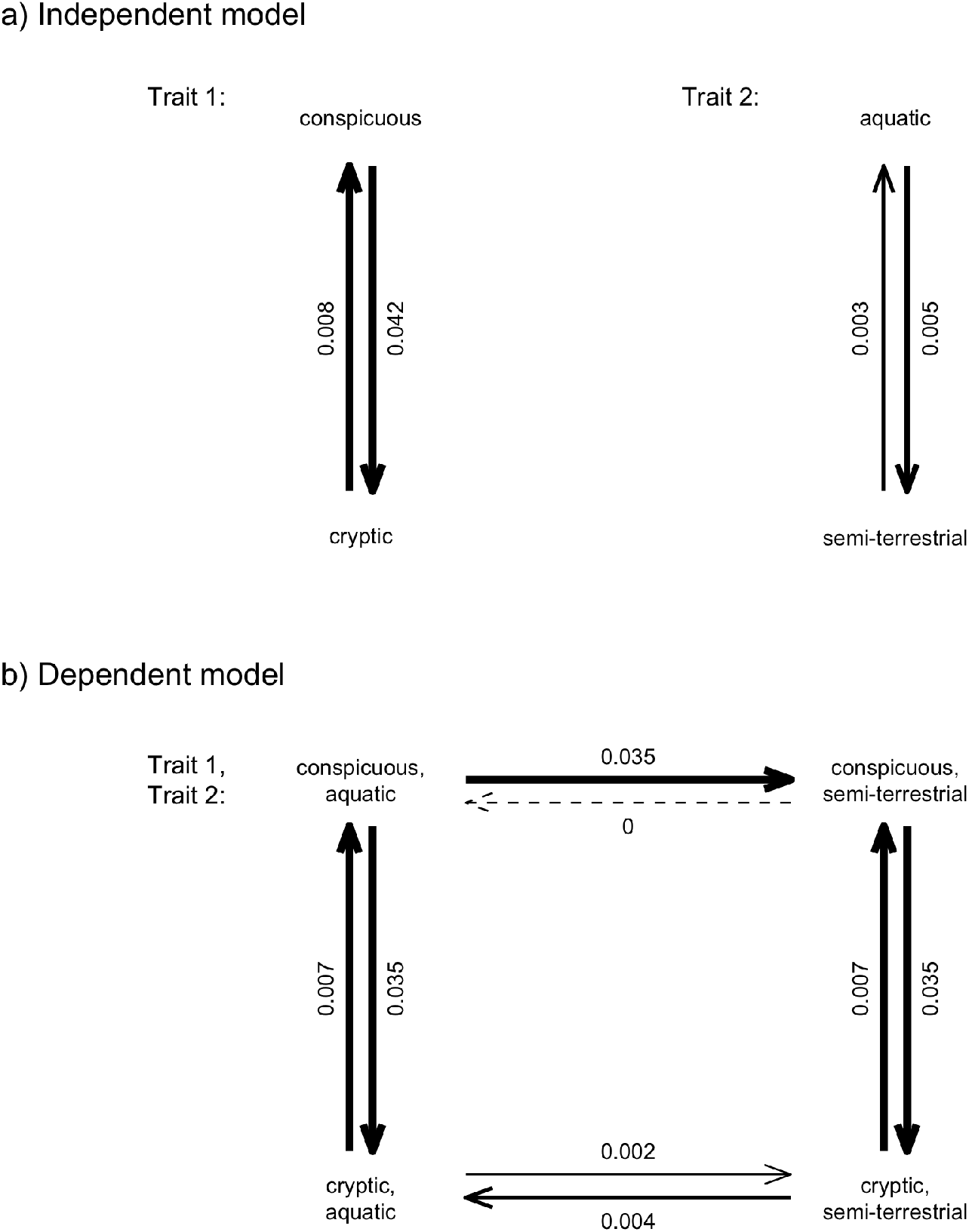
Fitted Pagel (1994) binary character evolution models for conspicuous coloration and burrowing in freshwater crayfish. Thicker arrows indicate higher transition rates.

A strong inference of correlated evolution between conspicuous coloration and burrowing behavior was supported, as the fitted parameters of the model implied that certain character combinations accumulated disproportionally over time when compared to other character combinations. For instance, we found that the rate of transition from conspicuous coloration and aquatic burrowing to conspicuous coloration and semi-terrestrial burrowing was relatively high. By contrast, the rate of transition from conspicuous coloration and semi-terrestrial burrowing to conspicuous coloration and aquatic burrowing was zero (Figure 3b). This observation suggests that if a lineage with conspicuous coloration were to evolve aquatic burrowing, our data and the fitted model suggest that conspicuous coloration would be lost likely instantly. Similarly, we found a relatively high transition rate from cryptic coloration and semi-terrestrial burrowing to cryptic coloration and aquatic burrowing, indicating that cryptic coloration has mostly prevailed in aquatic burrowers (Figure 3b). Within semi-terrestrial burrowers and aquatic burrowers, however, the transition rates from conspicuous coloration to cryptic coloration, and vice versa, appears to be equal.

## Discussion

The evolution and function of conspicuous coloration throughout semi-terrestrial burrowing crayfishes presents an evolutionary paradox to biologists; because these species rarely wander outside of their burrow and are nocturnal (Palaoro et al., 2013; Bearden et al., 2021). Despite this paradox, our work shows that conspicuous coloration has evolved over 50 times within the taxa we analyzed (*≈* 60% of described crayfish diversity). Blue, red, and orange coloration were the most common, having each evolved at least 15 times independently; whereas purple was the rarest color, having only evolved three times. Moreover, at least 10 species were coded as being polymorphic, with these species exhibiting multiple different phenotypes that are conspicuous (i.e., *Cambarus dubius*, *Euastacus sulcatus*). Clearly, in crayfish, conspicuous coloration is an evolutionary labile trait, with bright colors being taxonomically widespread throughout the phylogeny. The two most specious families (Cambaridae and Parastacidae) contain crayfish with all conspicuous coloration that we coded in our study, whereas Astacidae remains cryptic aside from one subspecies (orange in *Pacifastacus leniusculus klamathensis*). The taxonomically limited family Cambaroididae contained no taxa that were conspicuously colored. Despite being relatively common and apparently taxonomically widespread, evidence supporting the function (or lack thereof) of these colors are limited, although we believe that our correlated analysis of coloration and burrowing strategy yields insight into this trait’s evolution.

We found that conspicuous coloration in crayfish is evolutionary correlated to a semi-terrestrial burrowing strategy. Therefore, the typical association between light environment and adaptive coloration lacks support based on semi-terrestrial burrowing crayfish’s reclusive lifestyle; because semi-terrestrial burrowing species are nocturnal and based on current knowledge, rarely interact with each other on the surface (Diehl et al., 2022). Thus, major questions remain unresolved, for example: why would conspicuous colors be more common in species that rarely leave their burrow and are nocturnal?. Furthermore, are there costs and benefits to these conspicuous colors? We believe that recent developments regarding the evolution of neutral color phenotypes, that is, color phenotypes that may be under little to no selective pressures (Caro, 2021) are crucial in answering these questions. Several lines of evidence from disparate fields of research are needed to explain the link between coloration and semi-terrestrial burrowers. First, color mutations in crayfish and other decapods. Second, genetic and physicochemical underpinnings of crustacean coloration. And third, differences in natural history and gene flow of aquatic and semi-terrestrial burrowing crayfish. We explain these three lines of evidence below, which when considered together, suggests that conspicuous coloration in semi-terrestrial crayfishes might be a neutral color trait.

First, color phenotype mutations in crayfish and other decapods appear to be relatively common (Graham, 2023). Using mutant blue color phenotypes as an example, there are dozens of records from the primary literature of crayfish collected with a presumed mutation which results in an otherwise cryptic species expressing a conspicuous blue phenotype (Momot and Gall, 1971; Hayes II, 1975; Fitzpatrick, 1987; Secker, 2013; Kale et al., 2021). Similarly, in non-crayfish decapods, there are reports of blue clawed-lobsters (Christensson et al., 2013), spiny lobsters (Sampaio, 2021), and hermit crabs (Graham, 2023). Although no study has pinpointed the exact genetic mechanism responsible for these phenotypes, physiochemical evidence suggests that these phenotypes are a result of an overexpression of the blue crustacean-specific carotenoprotein *α−*crustacyanin (Cianci et al., 2002; Begum et al., 2015), which would result in the expected blue phenotype. Aside from the mutant blue phenotypes, crayfish have also been reported with other mutant color phenotypes such as red, orange, white, and others (Graham, 2023). Since chance color mutations appear to be common in decapods, these mutations are likely to control a wide range of color phenotype variation seen in crayfish. Similar mutation-based mechanisms are thought to control a wide range of color phenotypes in other animals (Bennett and Lamoreux, 2003; Braasch et al., 2007; Gehara et al., 2013; Stuckert et al., 2019).

Second, although the genes causing crayfish color mutations are unknown, at least in some species, mendelian inheritance studies have been conducted. For example, in both *Cherax destructor* and *Procambarus alleni* (two species with a cryptically colored wild type phenotypes), mutant blue phenotypes have been cultured. In both species, breeding experiments concluded that the mutant blue phenotype is derived from a single autosomal recessive mutation, because 100% of offspring produced from two individuals with the homozygous recessive phenotype express blue coloration (Black, 1975; Walker et al., 2000). Therefore, once two mutants arise in a population, blue color phenotypes may quickly become the dominant phenotype depending on the population size. Indeed, the simplicity of producing mutant blue crayfish phenotypes are utilized for commercial industries like the ornamental crayfish trade and aquaculture. In theory, although both species described above are taxonomically distant from one another, it is possible for the same mutation to appear independently (Gross and Wilkens, 2013; Martin and Orgogozo, 2013). However, the above two mechanisms (i.e., crustacean color mutations and the simple inheritance mechanisms of color mutations), must be combined with knowledge on the natural history and burrowing behavior of crayfishes to fully explain why semi-terrestrial crayfishes are more likely to be conspicuously colored.

A third line of evidence required to understand why we believe that conspicuous coloration serves as a neutral trait, requires a deeper understanding of crayfish burrowing strategies. Aquatic burrowing crayfishes are tied to lentic and lotic systems that are typically connected throughout a hydrological system (Hobbs Jr, 1981). In these species, gene flow and geographic distributions are high, as species are capable of migrating throughout their aquatic habitats (Wilson et al., 2004; Bubb et al., 2006). By contrast, in semi-terrestrial burrowing species, because these species are not tied to surface water (Hobbs Jr, 1981), they colonize semi-terrestrial habitats which are amenable to their burrowing lifestyle. The habitat requirements of semi-terrestrial burrowers are usually narrow, with many species only being collected within a fine range of ecological parameters (Loughman et al., 2012; Helms et al., 2013; Rhoden et al., 2016; Bloomer and Taylor, 2022). This leads to semi-terrestrial crayfish often occurring in small populations across highly disjunct populations (Shull et al., 2005; Schultz et al., 2009; Stern et al., 2017; Hurt et al., 2019). The habitats in which semiterrestrial crayfish live also limit their dispersal capabilities, with these species having expected low degree of gene flow and genetic distance across populations (Ponniah and Hughes, 2004; Shull et al., 2005; Hurt et al., 2019). As such, vicariance is thought to be a primary driver of speciation in semi-terrestrial crayfish species, as geographic barriers like mountains or rivers severely limited gene flow across populations (Thoma et al., 2023). In summary, aquatic-burrowing species have substantial opportunity for migration and gene flow between populations, whereas migration and gene flow in semi-terrestrial crayfishes are severely limited.

Combining our three lines of evidence, we can speculate on a scenario where a genetic mutation arises in a crayfish resulting in a mutant conspicuously colored phenotype. The genetic mechanisms controlling the mutant phenotype may be simple, such as a single autosomal recessive mutation (Black, 1975; Walker et al., 2000). If these two requirements occur in an aquatic burrowing species, selective pressure from visually guided predators would be predicted to select against this mutant phenotype. Even in scenarios with a lack of predators, a high degree of gene flow within these species would result in conspicuous phenotype being diluted by the wild type cryptic phenotype. However, in a semi-terrestrial species which rarely leave their burrows and has lower capabilities for dispersal and breeding with other populations, this color mutation might easily become widespread. Therefore, if a conspicuous color mutant arises in a semi-terrestrial species, we believe that it would be more likely to take hold due to low gene flow and less interaction with potential visually guided predators. We believe that these lines of evidence are the best explanation as to why semi-terrestrial burrowing crayfishes would be more likely to be conspicuously colored. In theory, mutations could arise within the same loci across the crayfish phylogeny, explaining the repeated evolutions of the same conspicuous colors we observed (red, orange, and blue). This model of repeated phenotypes across a clade has been demonstrated to be a widespread phenomenon in explaining phenotypic evolution throughout animals (Gross and Wilkens, 2013; Martin and Orgogozo, 2013). Indeed, similar mechanisms have been suggested to explain the conspicuous color polymorphism in coconut crabs (Caro, 2021). We acknowledge the high degree of speculation presented above, but we believe this type of provocative thought will be critical in furthering our understanding of the evolution of color phenotypes.

Although we support the notion that conspicuous coloration may evolve initially as a neutral color trait, it would be ignorant to conclude that conspicuous coloration is a neutral trait without empirical support (Caro, 2021). Unfortunately, there is poor evidence supporting the functional significance of conspicuous coloration in crayfish. We believe that the two primary selection pressures in other conspicuous animals, warning coloration and sexual selection, are unlikely to be strong selective pressures in this group. Regarding warning coloration, crayfish are unknown to possess any toxin, and typically resort to aggression when provoked by a predator (Robinson et al., 1970; Stein, 1976). Some crayfish species have claws which are differently or conspicuously colored than a typically cryptic body (i.e., *Faxonius virilis*, *Euastacus armatus*), which suggests that this color may serve as an aggressive warning signal to predators, but empirical tests of this hypothesis are lacking.

Regarding sexual selection driving conspicuous coloration in crayfish, color may function a signal of quality in male-male competition or female mate choice (Gherardi and Aquiloni, 2011). Interestingly, in the only study to date which quantified objective measurements of crayfish coloration, *Pacifistascus leniusculus*, it was found that there was sexual dimorphism in the brightness and saturation of male and female claws (Sacchi et al., 2021). Although no behavioral evidence was provided, these results suggest that there may be some underlying function with sexual selection in this species with conspicuous coloration, although further evidence is required to support these claims. Collection of objective color data with spectrometers is an obvious next step to further the hypothesis that coloration in crayfish serves a function in sexual selection.

Importantly, physiological evidence suggests that crayfish can detect at least some of the variation in coloration we describe herein. Some studies find that crayfish have a single-color photoreceptor, suggesting that they are capable of processing wavelengths of light in the red spectrum (Kennedy and Bruno, 1961). Alternative studies suggest that crayfish are dichromatic, which suggests that crayfish can detect both red and blue coloration (Wald, 1968, 1967; Suzuki and Eguchi, 1987; Crandall and Cronin, 1997). Interestingly, we found that blue, red, and orange were the colors most often exhibited by crayfishes; which overlaps with the range of colors crayfish are expected to detect. Overall, behavioral evidence is required prior to concluding any potential function of the conspicuous color diversity we described herein. Additionally hypothesis should also be explored, such as camouflage, heat absorption, or species identification (Detto et al., 2006; Schuster, 2020), all of which have been shown to provide a benefit to other conspicuously colored taxa. However, similar to coconut crabs, the nocturnality and lack of ultraviolet radiation suggests that carapace pigmentation is not used for heat management or UV protection semi-terrestrial burrowing crayfishes.

In conclusion, our work brings into question to traditional view of animal coloration as a perfectly adapted phenotype. The hypothesis that phenotypic traits such as coloration can evolve and persist as neutrally selected traits is not a novel idea (Ho et al., 2017), but finding natural systems to test such hypothesis are difficult. Across animals, the overly adaptations point of view that all coloration may be perfectly adapted or fit for the needs of the animal is not always expected (Caro, 2021). To progress our understanding of the neutral theory of color evolution, proximate mechanisms must be explored. Specifically, the genetic architecture of color is widely studied (Hoekstra et al., 2006; Braasch et al., 2007), but surprisingly little research has begun on this topic within decapod crustaceans. Crayfishes may serve as an ideal taxa for such genetic investigations, especially based on the knowledge that the genetic control of at least some color phenotypes are relatively simple (Black, 1975; Walker et al., 2000).

## Acknowledgements

N/A

## Data Accessibility Statement

A fully reproducible workflow of the data analyses, including R scripts and additional supporting material, is available in the following repositories: Github https://dylan-padilla.github.io/crayfish-coloration/, a Dryad link will be available upon acceptance.

## Author Contributions

Z.A.G conceived the study and collected the data; D.J.P.P analyzed the data and produced figures. Both authors drafted the manuscript, edited the manuscript, and agree to be held accountable for the content herein.

